# The Role of Rhizosphere Microorganisms and CNPS Genes in Shaping Nutritional Traits of Capsicum

**DOI:** 10.1101/2024.01.17.575995

**Authors:** Yu Tao, Mingxing Zhang, Siwen Peng, Shiping Long, Xuexiao Zou, Xin Li

## Abstract

The rhizosphere microbiota plays crucial roles in biogeochemical cycling and primary production. However, there is a lack of research exploring the complex relationships between microbiota and their functional traits in pepper rhizospheres, as well as their impact on nutrient cycling processes. Here, we investigated the effects of pepper species on the rhizomicrobiota and functional genes (C/N/P/S) on nutrient absorptions and accumulations in pepper organs. The results revealed that Pepper YZ/BE had higher N content in all compartments, which could be attributed to the presence of enriched N-metabolic microbes (*Gaiellales*/*Leifsonia*) and higher expression of N availability-promoting genes (*ureC*/*amoA2*/*nxrA*/*napA*) in rhizospheres. Additionally, we utilized co-occurrence network analysis and partial least squares path modeling (PLS-PM) to understand the interactions among the variables. The bacterial network exhibited more associations than the fungal network, and the abundance of certain modules positively correlated with the expression of CNPS genes, which thus significantly influenced pepper nutrient content. The PLS-PM analysis demonstrated that taxa abundance in network modules, functional genes, and rhizospheric soil properties collectively explained 92% of the variance in pepper nutrient content. Overall, this study provides valuable experimental and theoretical insights into the effects of rhizosphere microorganisms and CNPS genes on the nutritional traits of *Capsicum*.

**Highlight:** The rizho-bacterial community harbored more robust relationships than the fungal ones, which formed the functional clusters highly linking to the below- and aboveground nutrient properties of pepper species.

## Introduction

Plants have co-evolved with microbial communities for over 400 million years, resulting in highly diverse and specific taxonomic structures that colonize various plant tissues (Santoyo, 2022). Among these microbial communities, the rhizomicrobiota plays a crucial role in the soil-plant ecosystem by driving soil organic matter and nutrient transformation. They contribute to material circulation and metabolism, improve soil properties, regulate nutrient storage and release, and influence plant growth (Trivedi *et al*., 2020).

Numerous studies focusing on a wide range of crops have demonstrated a close relationship between soil rhizosphere microbial diversity and soil nutrient utilization (Berg, 2009; Liu *et al*., 2022; Perez-Izquierdo *et al*., 2019). These beneficial microbes, such as *Bacillus*, *Pseudomonas*, *Rhizobium*, *Trichoderma harzianum*, and *Aspergillus* spp., have been shown to promote plant growth, enhance nutrient absorption, and improve stress tolerance (Kour *et al*., 2021). Plant growth-promoting rhizobacteria (PGPR) and arbuscular mycorrhizal fungi (AMF) are two important groups of root-resident microbes that form functional clusters (Averill *et al*., 2019; Gouda *et al*., 2018). They enhance mineral bioavailability, secrete secondary metabolites like organic acids, poly-sugars, and amino acids, and help balance phytohormone levels, thus enhancing plant resistance to pathogens (Berlanga-Clavero *et al*., 2022; Gouda *et al*., 2018). Furthermore, rhizobacteria, particularly Bacillus spp., have been found to promote lateral root development (Li *et al*., 2021). The symbiotic microbiota that colonizes root zones also supports plant mineral nutrient homeostasis (Oldroyd and Leyser, 2020; Salas-Gonzalez *et al*., 2021). These findings highlight the dominant role of the rhizosphere microbial community and provide a solid foundation for further research in plant-microbiome breeding efforts.

The rhizosphere microbial community plays a vital role in promoting plant growth, nutrient absorption, and stress tolerance (Berg, 2009; Gouda *et al*., 2018). This symbiotic relationship between plants and microorganisms is reciprocal, as plants selectively influence the composition of the rhizosphere microorganisms to obtain specific functional traits necessary for their adaptability (Jiang *et al*., 2022; Iannucci *et al*., 2021). Through root secretions, plant hosts assemble a species-specific root microbiota that enhances nutrient solubilization and mobilization, leading to improved resource acquisition (Iannucci *et al*., 2021; Santoyo, 2022). This, in turn, affects the relative abundance of microbial taxa and reshapes the microbial community structure in the rhizosphere (Trivedi *et al*., 2020). The rhizosphere microbiome also secretes plant hormones that positively impact crop yield, quality, and soil nutrient availability (Santoyo, 2022). Furthermore, the unique rhizosphere microbial communities associated with different plant species contribute to variations in nutritional characteristics and crop quality (Bouffaud *et al*., 2014; Brisson *et al*., 2019). For instance, studies have shown that the absorption of nitrogen cycling-related bacteria contributes to differences in nitrogen use efficiency among rice varieties (Zhang *et al*., 2019). Similarly, the dissimilarity in microbial communities and plant genetic distances are highly correlated in canola and maize crops (Bouffaud *et al*., 2014). These findings highlight the intricate relationship between plants and the rhizosphere microbial community, emphasizing the importance of understanding and harnessing this relationship for agricultural advancements (Taye *et al*., 2019).

Rhizosphere microbial diversity is influenced by various factors such as plant development stages, ecotypes, genotypes within species, and soil environmental conditions like soil types, temperature, and moisture. The metabolic activity of rhizosphere microorganisms, specifically the expression of C/N/P/S genes, plays a crucial role in plant growth, nutrient uptake, and soil nutrient conditions. Previous studies in different crops including Arabidopsis (Li *et al*., 2021), rice (Wang *et al*., 2022), canola (Taye *et al*., 2019), and maize (Wagner *et al*., 2021) have demonstrated the significance of rhizosphere microbes in regulating soil nutritional conditions through their metabolic activities (Zhao *et al*., 2019). Additionally, functional microbes have been found to optimize nitrogen (N) and phosphorus (P) capture, thereby promoting overall plant growth (Oldroyd and Leyser, 2020). Strain-specific induction of genes involved in nutrient transformation and growth promotion has also been observed in growth-promoting strains (Wang *et al*., 2022). Large-scale investigations have shown positive correlations between specific microbial taxa and functional genes related to carbon (C) and nitrogen (N) cycling, as well as their associations with soil properties (Shi *et al*., 2020). Furthermore, differences in rhizosphere microbial composition and their effects on soil nutrient availability and aboveground traits have been observed among genotypes of the same species. For example, maize hybrids exhibit distinct rhizosphere microbiomes compared to inbred lines, and the heterosis of root and aboveground traits is strongly influenced by the belowground microbial community (Wagner *et al*., 2021). Similarly, different genotypes of pepper plants have been found to influence the physical-chemical and microbial properties of the surrounding soil (Khenaka *et al*., 2019). These findings highlight the important role of specialized microbial communities in improving the healthy development of plants by exerting their unique functional characteristics.

Capsicum annuum, a widely cultivated vegetable crop and condiment, belongs to the Solanaceae family. It is native to South America and Mexico and has been domesticated and grown globally. Despite the extensive diversity in fruit types and nutritional quality, the intricate relationship between the rhizospheric microbial community and the absorption of nutrients by pepper tissues remains poorly understood. This study utilized high-throughput sequencing technology to analyze the microbial diversity in the rhizosphere soil before and after pepper planting. It also examined gene expression related to nutrient transformation and absorption in the rhizosphere soil. The aim was to investigate the connection between nutrient accumulation in different pepper varieties and the diversity and metabolic activity of rhizosphere microorganisms.

## Materials and methods

### Experimental design

Experimental materials consisted of different pepper cultivars, namely yuanzhu (YZ), bell (BE), line (LN), and horn (HR), which exhibited distinct fruit-type variations. Prior to the experiment, it was observed that YZ and BE species had a higher capacity for nitrogen (N) absorption. Seeds of these cultivars were used to grow seedlings in a greenhouse under controlled conditions (30L, 60% humidity). After approximately 30 days, when the seedlings had developed around 8 leaves, they were transplanted into plastic pots (15:20:20 cm) filled with a mixture of laterite soil (from unplanted areas) and a seedling-exclusive matrix in a 2:1 ratio. The initial chemical composition of the mixed soil (designated as S0) was as follows: pH 5.6, total nitrogen (TN) 1.21 g/kg, total phosphorus (TP) 0.50 g/kg, total potassium (TK) 21.07 g/kg, available nitrogen (AN) 99 mg/kg, available phosphorus (AP) 33 mg/kg, available potassium (AK) 282.67 mg/kg, and organic matter (OM) 31.51 g/kg. During the pot experiment conducted from April to July 2021, a general chemical fertilizer with a ratio of N:P:K = 17:17:17 was periodically applied to the soil along with watering to ensure the normal growth of the pepper plants. This practice was carried out at ten-day intervals, amounting to a total of six applications throughout the pepper’s entire growth cycle.

### Sample collection and determination

During the flowering stage, rhizosphere soils were collected from each pepper plant by gently shaking the roots. Additionally, four parts of the pepper plant, including the roots, stems, leaves, and fruits, were harvested. These plant parts were then dried at 80, crushed, and passed through a 2mm sieve to obtain fine powder samples. The main nutrient content of these samples, such as total nitrogen (TN), total phosphorus (TP), and total potassium (TK), was determined. The measurement of total nutrient content in the rhizosphere soil, as well as in the plant parts, and the available soil nutrients (AN, AP, AK) in the rhizosphere soil, were conducted following the methods outlined in the book “Soil Agrochemistry Analysis” by Bao (2005) (Bao, 2005).

### DNA sequencing analysis

Soil total DNA was extracted using the E.Z.N.A.® soil DNA Kit (Omega Bio-tek, Norcross, GA, U.S.) following the manufacturer’s protocols. The primer pairs 338f/806r (5’- ACTCCTACGGGAGGCAGCA-3’/5’-GGACTACHVGGGTWTCTAAT-3’) and ITS1f/ITS2r (5’- CTTGGTCATTTAGAGGAAGTAA-3’/5’-GCTGCGTTCTTCATCGATGC-3’) were used to amplify the V3-V4 region of the 16S rRNA gene and the ITS1 region of the rRNA gene, respectively. The extracted DNA was electrophoresed on a 1% agarose gel and checked with a NanoDrop 2000 UV-vis spectrophotometer (Thermo Scientific, Wilmington, USA). PCR reactions were performed as described in a previous paper (Shen *et al*., 2020). The initial PCR products were checked on a 2% agarose gel. The 16S rRNA amplicons and ITS amplicons were pooled separately and sequenced using the Illumina MiSeq instrument (Illumina, San Diego, CA, USA). The raw sequences were quality filtered using the Quantitative Insights into Microbial Ecology pipeline (QIIME v1.9), and taxonomy was assigned using the Ribosomal Data Project classifier at 97% sequence similarity to generate the OTU dataset. After filtering, there were 2,725,114 bacterial gene sequences and 3,258,085 fungal gene sequences, representing 2,000 bacterial OTUs and 547 fungal OTUs. The sequencing data has been uploaded to the NCBI Sequence Read Archive (SRA) database (accession number PRJNA981279)(https://submit.ncbi.nlm.nih.gov/).

### High-throughput quantitative PCR assay

The DNA extracted from soil samples was assessed for quality using the Qubit 4.0 instrument from Thermo Fisher Scientific. This study used 71 primers to examine genes related to CNPS-cycling, which included six function clades: carbon degradation, carbon fixation, methane metabolism, nitrogen cycling, phosphorous cycling, sulfur cycling, and a reference gene (16S rRNA) (Table S2, S3). The qPCR reaction and fluorescence signal detection were performed using the SmartChip Real-Time PCR System from WaferGen Biosystems USA. Canoco software analyzed the Ct values obtained for each gene in each sample, and the relative quantitative values of the genes in each sample were calculated using the formula: 10^(31-Ct)/(10/3). The relative expression of 71 individual genes was visualized using a heatmap generated by the R package pheatmap. Non-metric multidimensional scaling (NMDS) was used to determine the distribution differences among the soil samples from the pepper rhizospheres based on the Bray-Curtis distances. This methodology was previously described by Zhao *et al*. (2019) and Zheng *et al*. (2018) (Zhao *et al*., 2019; Zheng *et al*., 2018).

### Bioinformatic and statistical analysis

For the bioinformatic and statistical analysis, the ACE and Shannon indices were used to calculate the richness and diversity of the microbial community, respectively, at the OTU level. The R vegan library was utilized to perform this analysis on rarified data. To visualize the dissimilarity between samples, principal coordinates analysis (PCoA) was conducted based on bray-curtis distances. The relative abundance of the microbial community at the phylum and genus levels was aggregated and plotted using Origin 9.0.

To identify significant differences in taxa between pepper rhizospheric soils, the Galaxy web application (http://galaxy.biobakery.org/) was used to conduct linear discriminant effect size analysis (LefSe) with an LDA score of 4. The co-occurrence of bacteria and fungi communities in pepper cultivars was determined using Operational Taxonomic Units (OTUs) (Segata *et al*., 2011). A correlation matrix was created using the R package igraph by calculating pairwise Spearman correlations. A significant co-occurrence was defined as a statistically robust correlation between OTUs with an r > 0.65 and a P-value < 0.05, with P-values adjusted using FDR-controlling adjustment.

The co-occurrence network was constructed by selecting a specific number of microbes and resulted in 887 OTUs for the bacterial interaction network and 230 OTUs for the fungal interaction network. Nodes represented OTUs, and edges represented high and significant correlations between nodes. The Gephi 9.0 software was used for network visualization and modular analysis (Ken, 2015), with each module indicating potential interactions and habitat sharing (Menezes *et al*., 2015).

To examine the relationships between modules in the network and the characteristics of pepper nutrients, the Spearman method was used. The results were visualized in the form of a heatmap using the R package pheatmap (Gu and Hubschmann, 2022). Intra-correlation calculations were performed for the CNPS genes from rhizospheres, core modules in the network, soil characteristics of rhizospheric zones, and pepper nutrients in different nutritional organs (root, stem, leaf, fruit). Partial Least Squares Path Modeling (PLSPM) was conducted using the ’plspm’ package (Yang *et al*., 2017) to gain a comprehensive understanding of the effects of microbiota and their functional genes in pepper rhizospheric soils on pepper nutrient properties. The different effects of pepper cultivars on each tested item were determined using one-way analysis of variance (ANOVA) with the Turkey test, performed using SPSS 24.0 software.

## Results

### Nutrients comparison in rhizosphere and pepper parts

The pepper cultivars used in this experiment exhibited significant phenotypic differences, particularly in the performance of their leaves and fruits. We conducted sampling two and a half months after transplanting during the fasting growth stage of the peppers. We analyzed the basic nutrients (total nitrogen, phosphorus, and potassium) present in the peppers from both above and below ground, as shown in Fig. 1 and Table S1. In the rhizosphere, there were no significant differences in the soil properties of TN, TP, AP, TK, and pH, except for the available soil nutrients of AN and AK (Table. S1). The YZ and BE pepper species had relatively higher AN nutrients in the soil compared to the other two species. The TN contents of pepper parts in root, stem, and leaf changed in the same way, with YZ and BE species showing higher TN content, similar to TP in the parts of stem and fruit. However, TP in the root and leaf showed no significance among the peppers, nor in the rhizospheric soils. Furthermore, TK content in all the parts clearly changed among the four cultivars, which was not in line with the changes observed in the other parts’ nutrients of peppers (Fig. 1).

**Fig 1.**
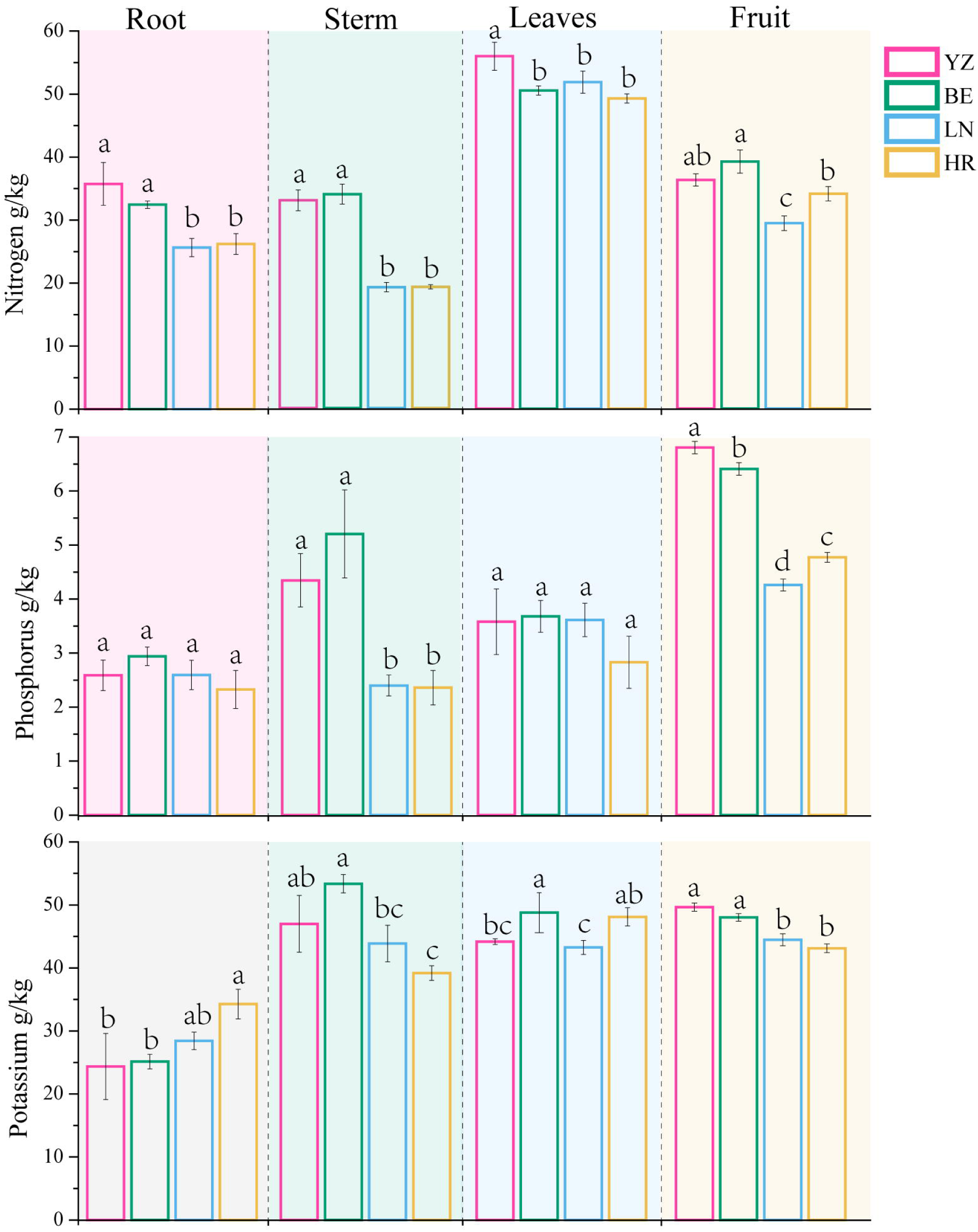
The basic nutrients content of nitrogen, phosphorus, potassium from pepper parts of root, stem, leaves, fruit.

### Microbial diversity and composition of rhizosphere soils from pepper cultivars

The bacterial community in the rhizosphere of pepper plants (YZ, BE, LN, HR) exhibited higher richness and diversity compared to the fungal groups (Fig. S1). The planting of peppers significantly increased the microbial community diversity in the rhizosphere soils, as indicated by the ACE and Shannon indices (Fig. S1 a-b). The Principal Coordinate Analysis (PCoA) further demonstrated the dissimilarity of microbial communities between the unplanted soil (S0) and the rhizosphere soils of pepper plants (Fig. S1 c-d). Notably, the ACE and Shannon indices, along with the PCoA analysis, revealed distinct differences between the YZ and BE pepper species and the other two species (Fig. S1a-c).

The relative abundance of bacterial phyla, such as Actinobacteriota, Firmicutes, and Proteobacteria, was predominant in both pre-and post-planting soils, accounting for up to 74%-85% of the total. However, after pepper planting, there was a significant decrease in Actinobacteriota, Firmicutes, as well as other phyla like Bacteroidota, while Proteobacteria, Patescibacteria, Acidobacteriota, and Gemmatimonadota showed an increase (Fig S4). LefSe analysis, tested by the Kruskall-Wallis test, revealed significant differences in microbial composition at the phylogenetic level, with certain phyla (YZ, BE, LN, HR) acting as biomarkers for pepper planting (Table S3). At the genus level, the most abundant genera in non-planted soils (S0), namely *Streptomyces* and *Rhodanobacter*, declined by over 50% after pepper planting, along with *Kribbella* and *Hydrogenispora* (Fig S4). On the other hand, genera like *Sinomonas*, *Bacillus*, *Arthrobacter*, *Bradyrhizobium*, *Leifsonia*, *Burkholderia*-*Caballeronia*-*Paraburkholderia*, *Sphingomonas*, and *Knoellia* increased in abundance in the pepper rhizosphere soils compared to S0. Among the pepper rhizosphere soils, there were significant differences in the relative abundance of genera among the soil samples, as summarized based on LDA scores (>4). The YZ species had 29 different genera, mainly belonging to Proteobacteria and Actinobacteriota, while the other species (BE, LN, HR) had 10, 10, and 20 different genera, respectively (Fig. 2, Table S4). It is worth noting that the pepper species YZ and BE dominated the genera *Gaiellales*(o), *Leifsonia*, *Conexibacter*, and *Frankiales*(o), whereas the other species (LN and HR) had enriched genera such as *Rhodanobacter*, *Rhodococcus*, *Saccharimonadales*(o), and *Steroidobacter*. In terms of fungi groups, the phylum Ascomycota was predominant in raw soil or rhizosphere soils of pepper, ranging from 62% to 94%. However, its abundance decreased in pepper planting soils, especially in YZ soil. Conversely, the phyla Basidiomycota and Rozellomycota increased in pepper soils. At the genus level, the relative abundance of *Chaetomidium*, *Cephalotrichum*, *Pseudogymnoascus*, and *Umbelopsis* significantly decreased in pepper rhizosphere soils compared to unplanted soil (S0), with *Chaetomidium* showing the largest decrease. On the other hand, the genera *Candida*, *Gibberella*, and *Rhodotorula* increased in abundance and filled the new ecological niche in the pepper rhizosphere (Fig. S2).

**Fig 2.**
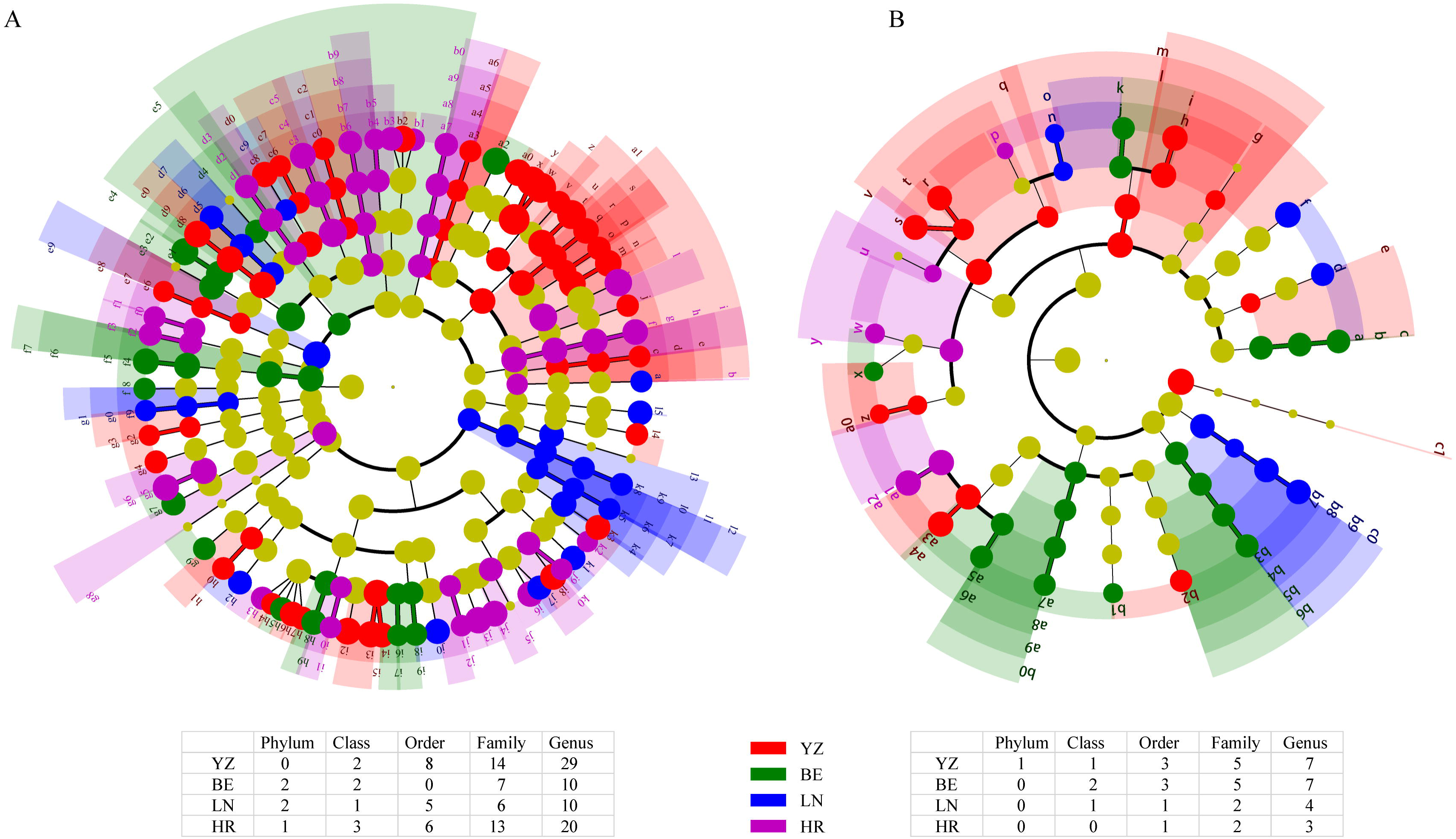
Linear discriminant analysis (LDA) effect size analysis (a, bacteria; b, fungi) of four pepper cultivars. LDA score > 4 indicates a biomarker with statistical differences between peppers. The letters (a-x) denote different classification levels from phylum, class, order, family and genus. Sunburst charts respectively colored by peppers show significant differences at *P* < 0.05 level. The tables list microbial biomarkers among four peppers at different taxonomic levels.

### Co-occurring network analysis

We then analyzed the co-occurring bacterial and fungal communities in pepper rhizosphere soils, representing them as network modules, which can be thought of as ecological clusters. For the bacterial community, we used 887 OTUs to create an interplay network at the OTU level, retaining 2418 statistically significant edges among 621 nodes (r<0.65 & *P*<0.05). The majority of these edges (88.84%) represented positive relationships within the network. In the bacterial community, the phyla Proteobacteria and Actinobacteriota were prominent in both pepper rhizosphere soils and the co-occurrence network (Fig. 3A). They were particularly abundant in three large modules: module#3 (110 OTUs), module#11 (124 OTUs), and module#8 (78 OTUs) (Fig. 3B, Table S5). Other modules exhibited different predominant bacterial phyla, such as Chloroflexi in module#13, Actinobacteriota in module#15, and Firmicutes in module#17. Notably, there were stronger interactions within the modules compared to between them, with module#3, module#8, and module#11 having 540, 433, and 580 edges, respectively, while there were relatively fewer associations between network modules. In contrast, the fungal network, constructed with only 230 OTUs using the same cutoff values, appeared less complex with fewer relevant edges, consisting of 112 edges among 85 nodes (Fig. S3). The Ascomycota phylum occupied a central position in the fungal network.

**Fig 3.**
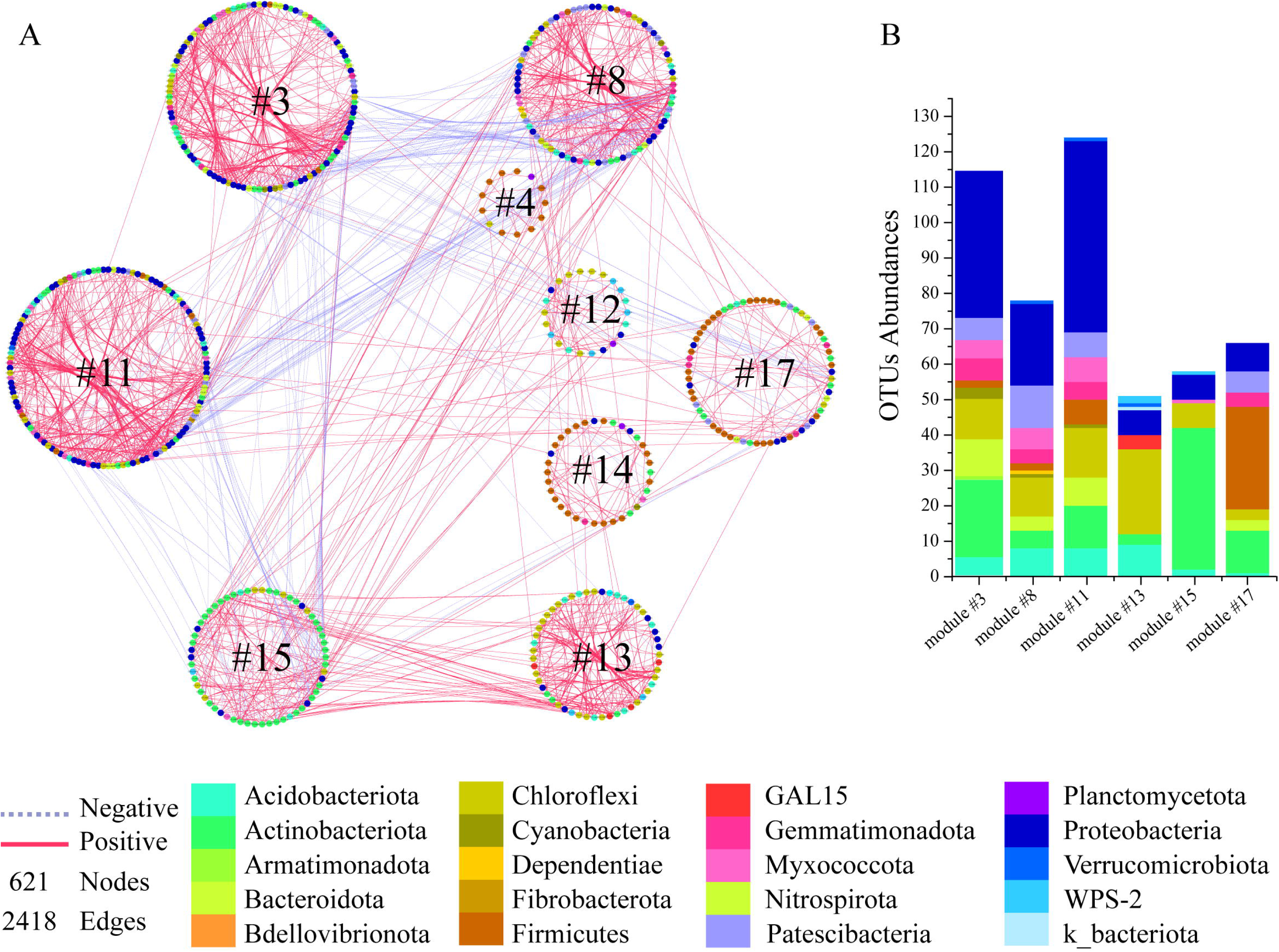
A. Co-occurrence network of rhizosphere soils of peppers with strong and significant correlations (|r| >0.65 & *p* <0.05). B. The abundances of OTUs belonging to network modules. The network is colored by phylum levels and clustered by modularity features. Nodes within the same modules represented a closely interconnected relationship.

Interesting findings were observed in the study. Certain bacterial genera, such as *Gaiellales*(o), *Leifsonia*, *Conexibacter*, *Frankiales*, and *Caulobacteraceae*, were found to be enriched in both YZ and BE species, specifically in module#15. On the other hand, genera like *Rhodanobacter*, *Rhodococcus*, *Saccharimonadales*, *Steroidobacter*, LWQ8, and *Bradyrhizobium* dominated in LN and HR, mostly in module#11 and module#17. Additionally, LDA analysis identified 16 key genera in YZ, including *Arthrobacter*, *Sinomonas*, *Knoellia*, and *Bacillus*, which were predominantly assembled in module#3. Similarly, module#11 contained 8 abundant bacterial taxa from HR. These findings suggest both differences and similarities between YZ&BE and LN&HR in terms of bacterial composition.

### Functional differentiation of rhizosphere microbes among pepper cultivars

The QMEC (Quantitative Microbial Elemental Cycling) method, utilizing HT-qPCR technology, was employed to comprehensively analyze the abundance and diversity of functional genes involved in carbon (C), nitrogen (N), phosphorus (P), and sulfur (S) cycling in soil samples from different pepper cultivars. Notably, certain genes such as *exo*-*chi*, *iso*-*plu*, *hzo*, and *cphy* were absent in all soil samples. The comparison of CNPS cycling gene profiles between raw soils and pepper rhizosphere soils, as indicated by NMDS analysis (PERMANOVA, *P* < 0.05), revealed significant differences (Fig. 4A). The expression levels of C fixation, N and P cycling genes were considerably higher in YZ cultivars, while the C/N metabolic genes were more abundant in BE and LN species, compared to raw soil and other rhizosphere soils (ANOVA, *P* < 0.05) (Fig. 4B). The HR cultivar exhibited the lowest abundance of functional genes. Alongside the variation in total gene expression, the 68 detected functional genes among soil samples also displayed some differences (Fig. 4C). Following planting, several genes related to N and S cycling, such as *nosZ1*, *nxrA*, *ureC*, *amoA2*, and *phoD*, showed a decline in expression levels in the rhizosphere soils compared to the raw bulk soil (S0). Among the pepper cultivars, YZ and BE rhizosphere soils exhibited a greater number of highly expressed genes, unlike LN and HR species. Notably, the genes *phoX*, *phoD*, and *phnK* (P cycling), *pmoA* (C methanol), *amyA* (C cycling), *lig* (C degradation), *NirS1*, *gdhA* (N cycling), and *cdaR*, *pccA*, *korA* (C fixation) showed high expression levels in YZ and BE cultivars, whereas few highly expressed genes were found in LN and HR rhizosphere soils. Genes such as *ureC*, *amoA2*, *hao*, *nifH*, *nasA*, *nxrA*, and *napA*, which are involved in N cycling and contribute to the availability of soil nitrogen, exhibited higher expression levels in YZ and BE cultivars compared to other cultivars (Fig. 4C).

**Fig 4.**
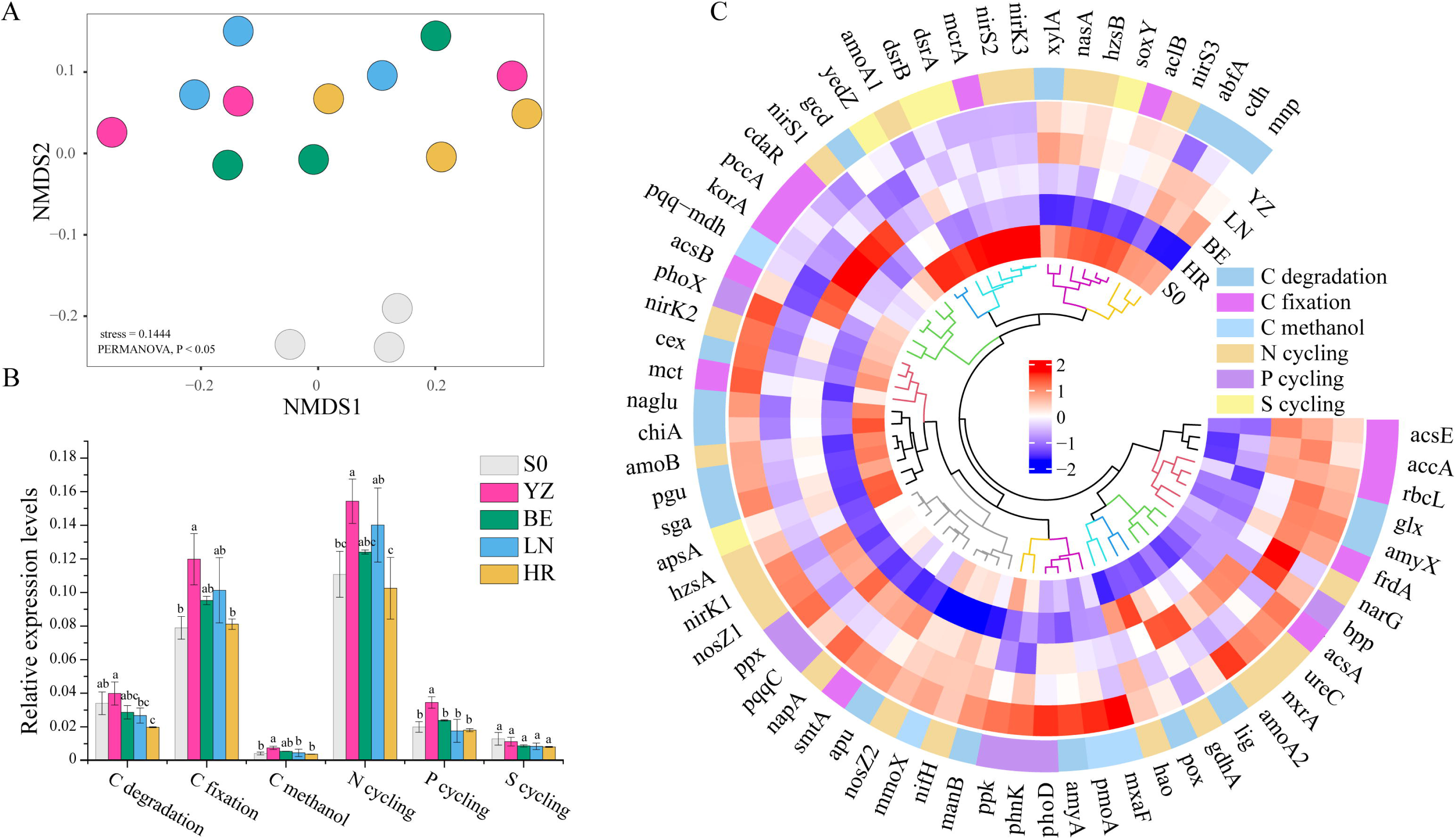
Non-metric multidimensional scaling (NMDS) based on the Bray Curtis distances reflecting the distinct distribution patterns of CNPS cycling gene profiles in all soil samples (A). The relative abundance of CNPS gene profiles in four pepper rhizosphere soils in form of subtotal groups (B) and of individual genes (C). There are 71 functional genes (except for 3 genes with no detection), which belong to six functional groups, such as C degradation (19), C fixation (13), C methanol (4), N cycling (22), P cycling (8), S cycling (5).

### Links between rhizosphere microbes and CNPS cycling genes

The study examined the correlation between microbial module groups and CNPS cycling functional genes in pepper rhizosphere soils using the Spearman correlation coefficient. Interestingly, three bacterial network modules (i.e., module#3, module#8, module#15) showed positive correlations with almost all gene groups, whereas module#11, module#13, module#17 exhibited completely opposite results (Fig. 5). Of particular note, module#11 displayed significant correlations with six functional genes related to C degradation, C fixation, C methanol, and N/P/S cycling at a p-value of less than 0.05. In contrast, module#11 showed 34 negative correlations with 67 genes, including 11 genes for C degradation, 6 for C fixation, and 13 for N/P cycling. However, more positive correlations were found in module#3, module#8, and module#15. Module#8 had the most positive relationships with genes belonging to C degradation, C fixation, and N/S cycling groups, with a total of 25 genes with significant levels (*P* < 0.05), followed by module#15 with a few similar genes.

**Fig 5.**
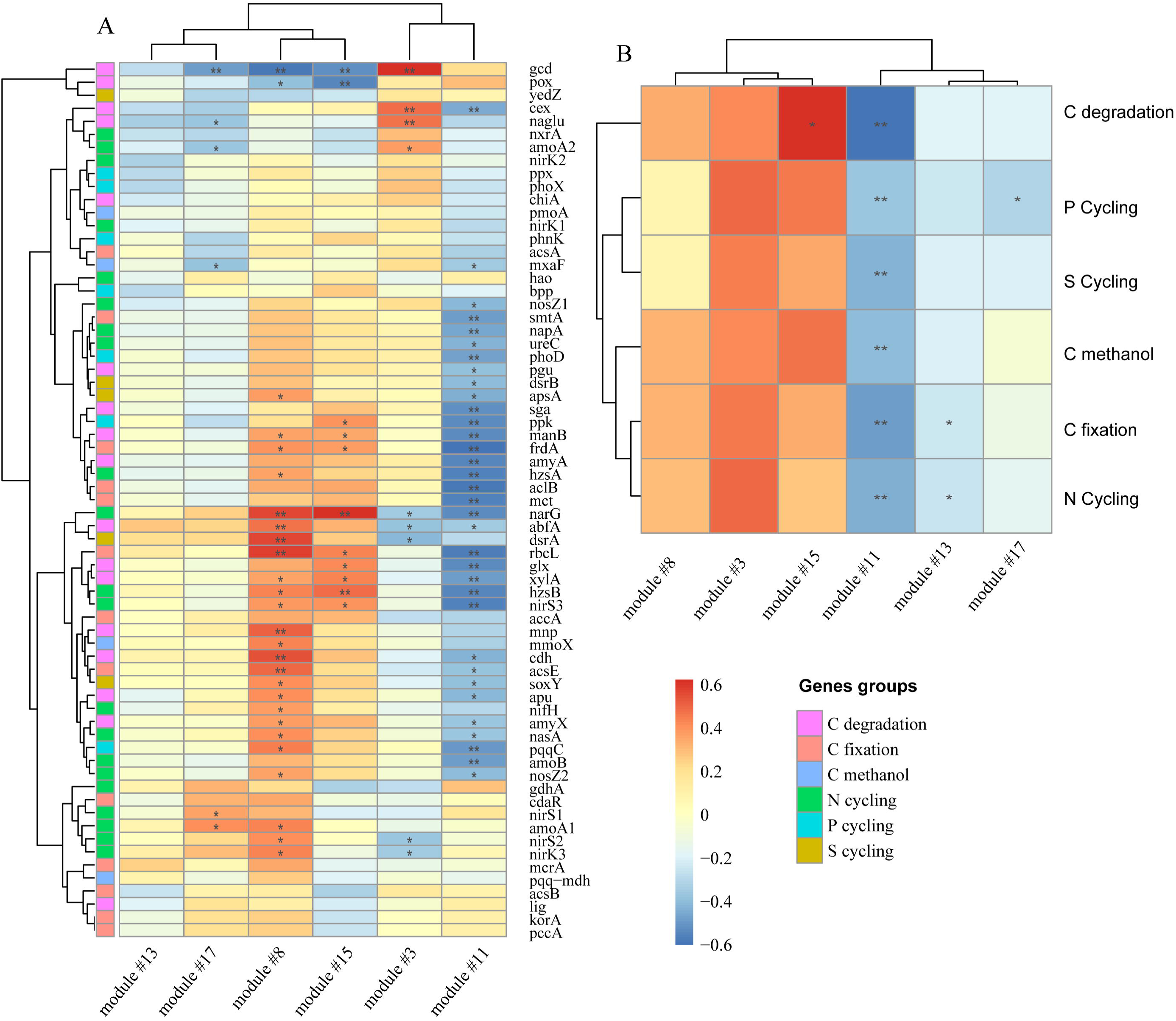
Correlation coefficients between the abundance of modules from bacterial network and the expression levels of CNSP cycling-related genes in pepper rhizosphere soils in the form of individual (A) or groups (B). Red cell represents for positive correlation and blue for negative ones. \*\**P* < 0.01; \**P* < 0.05.

### Effects of soil microbes, and the CNPS-related genes on pepper nutrients

In our study, we explored the correlations between functional genes of microbes in pepper rhizosphere soils and nutrient levels in the soils or different parts of the pepper plant. We found significant relationships among these variables. The soil properties such as pH and TK, as well as TK levels in pepper roots and leaves, showed negative but significant associations with functional gene groups. Conversely, changes in AN content in rhizosphere soils and the nutrient properties of pepper roots, stem, and leaves exhibited positive correlations with the expressions of the six functional genes (*P* < 0.05) (Fig. S4B). When considering individual genes, their significant correlations were mainly concentrated in three distinct zones on the heatmap (Fig. S4A). Furthermore, Partial Least Squares Path Modeling (PL-SPM) analyses revealed the intricate relationship between bacterial properties, functional genes, and pepper nutrient variables (Fig. 6). The CNPS-cycling genes strongly influenced soil physiochemical properties (R^2^=0.52) and significantly impacted root nutrient levels. Moreover, the nutrient properties of rhizosphere soils and roots had a substantial effect on aboveground nutrient levels in pepper plants, with the root’s effect being particularly pronounced. Ultimately, the latent variables explained a significant proportion of the variation in aboveground nutrient properties, collectively accounting for 92% of the variation.

**Fig. 6.**
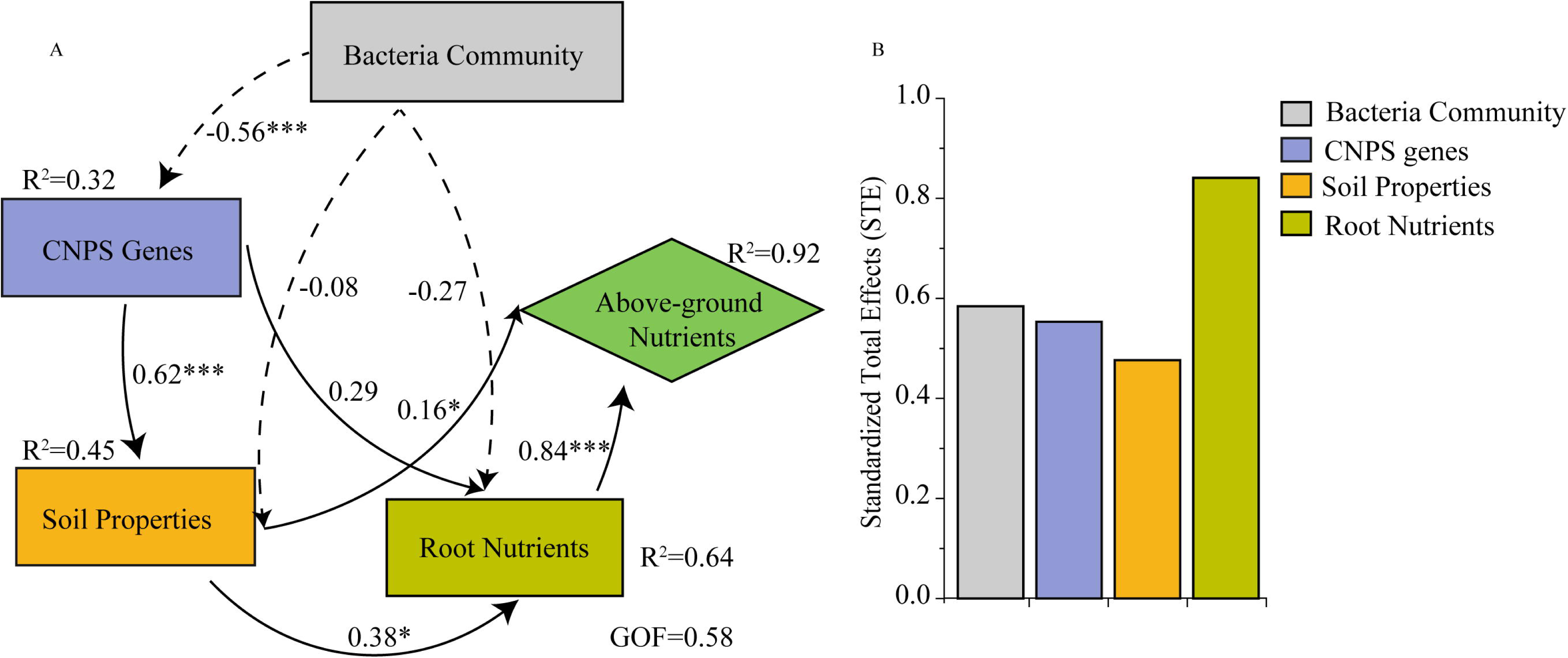
Partial Least Squares Path Modeling (PLS-PM) of the relationships among these latent variables (i.e., bacterial community, CNPS genes, soil properties, root nutrients, and above ground nutrients) (A); the standardized total effect on pepper above ground nutrients properties (B). Solid lines represent positive effects and broken lines represent negative effect. Numbers on the lines in the PLS-PM model are the path coefficient values (standardized effect sizes).

## Discussion

### Diversity of microbial community and occurrence network

The rhizosphere harbors complex microbial communities that play vital roles in various ecological processes (Fan *et al*., 2019; Shen *et al*., 2020). Our study revealed that the bacterial networks in the rhizosphere exhibited stronger interplay and robustness compared to the fungal networks. These bacterial networks formed distinct clusters, indicating their importance in nutrient transformation, nitrogen fixation, and the regulation of soil and climate variables (Fan *et al*., 2019; Zhang *et al*., 2018).

Manipulating these microbial communities holds great potential for enhancing agricultural practices, leading to improved crop productivity and more sustainable farming systems. The rhizosphere, the region surrounding plant roots, was found to host a higher microbial diversity and community composition compared to the bulk soil (Fan *et al*., 2017). Our research demonstrated that the presence of host plants significantly influenced the composition of the rhizosphere microbial community, as observed after pepper planting. We conducted a Kruskall-Wallis test and observed significant differences in rhizospheric microbial taxa among the four pepper species (Fig. 2). Interestingly, these differences were more pronounced in the bacterial community compared to the fungal community. We identified 29, 10, 10, and 20 different genera of bacteria in the four pepper species, while there were only 7, 7, 4, and 3 different genera of fungi. This indicates that rhizobacteria, bacteria that inhabit the rhizosphere, are more susceptible or active in the experimental soil conditions compared to fungi (Zhu *et al*., 2021). Furthermore, we performed PCoA analysis to assess the dissimilarity of the microbial community structure among the four pepper species. The results demonstrated that the genotype of the peppers significantly influenced the microbial community composition, consistent with previous findings in other crops (Bouffaud *et al*., 2014; Brisson *et al*., 2019). Specifically, we observed that the bacterial and fungal community compositions of YZ&BE peppers were more similar to each other compared to the other peppers. Conversely, HR&LN peppers exhibited a distinct community composition. These findings suggest the presence of specific microbial associations and interactions influenced by the genetic makeup of the pepper varieties.

At the genus level, the rhizosphere of YZ&BE peppers was found to be abundant in genera such as *Streptomyces*, *Bacillus*, *Leifsonia*, *Frankiales*(o), *Gaiellales*(o), *67-14*(f), and *Nocardioides*. These genera are known for their important roles in plant nutrition and growth. For instance, Streptomyces and Bacillus have been identified as core hubs in correlative networks and potential antagonists to Fusarium wilt disease (Gao *et al*., 2021). They are also involved in nutrient cycling, such as nitrogen fixation by *Frankiales*(o), *Streptomyces*, *Bacillus*, and *Arthrobacter* (Sellstedt and Richau, 2013), nitrate-nitrogen improvement by *Nocardioides* (Gao *et al*., 2022), and denitrification by *Bradyrhizobium* and *Mesorhizobium* (Zhang *et al*., 2023). These findings highlight the complex interactions within the rhizosphere microbial community and their impact on plant health and nutrient availability.

The rhizosphere, which is the region surrounding plant roots, is a dynamic interface where plant roots interact with the soil. It is home to a diverse community of microbes that play crucial roles in nutrient cycling and plant health. In this study, we focused on examining the microbial networks in the rhizosphere of different pepper varieties. We found that the bacterial network exhibited greater robustness and interconnectedness compared to the fungal network (Trivedi *et al*., 2020). This suggests that bacteria have a higher degree of interaction and cooperation in processes related to nutrient cycling. By analyzing the network modules, we identified specific bacterial groups that were highly connected within the network. These groups included Actinobacteria, Proteobacteria, and Chloroflexi, which are commonly found in healthy soils and have been associated with various beneficial functions, such as nitrogen fixation and nutrient transformation. Previous studies have also emphasized the significance of bacterial network modules in agricultural management. For example, a specific module (#3) associated with nitrogen fixation was found to decrease under long-term fertilization practices, indicating its importance in sustaining nitrogen fixation rates (Fan *et al*., 2019). Furthermore, the modules observed in both bulk and rhizospheric soils showed positive correlations with edaphic (soil) and climatic variables. This suggests that these modules are involved in nutrient transformation and regulation processes (Zhang *et al*., 2018), influenced by soil and climatic conditions. These findings highlight the intricate relationships between the rhizosphere microbial community and its surrounding environment, emphasizing the importance of understanding these interactions for sustainable agriculture and plant health.

### Microbial Modules and Ecological Functions

The composition and interactions of microbial communities in the rhizosphere have a significant impact on the nutrient cycling process. The microbial network in the rhizosphere forms complex ecological clusters that play a crucial role in nutrient transformation and agricultural management. These bacterial network modules are taxonomically diverse and can strongly correlate with the changes of C/N/P/S cycling genes.For instance, module#15, which includes microbes from phyla Actinobacteriota, Chloroflexi, and Proteobacteria, has been found to have a significant correlation with the abundance of 7 C metabolism-involving genes and 3 N cycling genes. This indicates that the microbes in this module may play important roles in C degradation and fixation, as well as N cycling. The Actinobacteriota phylum, in particular, has the potential to decompose xylogen and cellulose (Eriksson *et al*., 1990; Liu *et al*., 2020). Additionally, other bacterial families involved in N cycling, such as *acetobacteraceae* (Proteobacteria phylum) (Qiu *et al*., 2021; Wolfe *et al*., 2014), are also present in module#15.

Further analysis revealed significant positive correlations between module#8 and module#13 genes involved in carbon cycling (*abfA*, *cdh*, *mnp*, *acsE*, *rbcL*, etc.), as well as 10 genes related to nitrogen cycling and sulfur cycling genes (*apsA*, *dsrA*, *soxY*). The microbes associated with module#8 were classified into the phyla Acidobacteriota, Actinobacteriota, Bacteroidota, Gemmatimonadota, Patescibacteria, and Proteobacteria, all of which have been extensively studied for their roles in C/N/P/S cycling. For example, a study by Lan *et al*. (2022) demonstrated that changes in bacterial taxa from Acidobacteriota, Actinobacteriota, and Proteobacteria due to afforestation significantly influenced soil organic carbon fractions (Lan *et al*., 2022). Furthermore, Acidobacteria, Bacteroidetes, and Actinobacteria have been found to efficiently decompose plant and fungal cell wall biopolymers (Tlaskal and Baldrian, 2021), while Proteobacteria are known for their rich gene complements involved in N cycling in deadwood.

Microbial communities have been found to play a critical role in nitrogen cycling, with specific groups of microbes involved in processes such as nitrogen fixation, nitrification, and denitrification. These N-cycling microbes exhibit diverse functions and phylogenies, including N2-fixing cyanobacteria, nitrite-oxidizing bacteria (primarily found in Proteobacteria and Chloroflexi phyla), and denitrifying bacteria (Proteobacteria groups) (Isobe and Ohte, 2014; Li *et al*., 2021). Interestingly, the genes associated with module#8 showed positive responses to environmental variables in the co-occurrence network while exhibiting negative responses to module#11. These two modules occupied different niches in the co-occurrence network, indicating distinct ecological functions (Fan *et al*., 2019; Jiao *et al*., 2022). Understanding the composition and interactions of microbial communities in the rhizosphere can provide valuable insights into optimizing agricultural practices and enhancing nutrient cycling for improved crop productivity.

### Effects of Rhizomicrobiota and their functional genes on pepper nutrient properties

Soil microbial communities play a crucial role in regulating the biochemistry cycles of carbon (C), nitrogen (N), phosphorus (P), and sulfur (S) in the soil, which are essential for plant growth and the overall health of the ecosystem. In the context of pepper rhizosphere soils, LefSe analysis revealed significant taxonomic differences among the microbial communities, leading to the specific recruitment of microbes in the surface layer of the root (Table S4). The nitrogen content varied among the different pepper samples, with variations in microbial composition suggesting differences in the nitrogen cycling processes between the different pepper species. The rhizospheric soils of YZ and BE cultivars showed a higher abundance of genes related to nitrogen cycling, contributing to the higher nitrogen content in these plants. Similarly, Jiang *et al*. (2017) and Pérez-Izquierdo *et al*. (2019) found that plant genotype has a profound effect on the phylogenetic structure of the root-associated microbiota, influencing carbon regulation and nitrogen cycles (Jiang *et al*., 2017; Perez-Izquierdo *et al*., 2019). Additionally, researchers have explored the relationships between root traits and microbial communities (Lau and Lennon, 2011; Spitzer *et al*., 2021), indicating that below-ground microbial communities play a vital role in shaping plant evolutionary processes, ultimately affecting the productivity, diversity, and composition of plant communities (Wagg *et al*., 2011). In our study, we examined the differential expression of C/N/P/S genes in the associated microbes among different pepper cultivars, finding that these gene expressions were closely linked to the accumulation of below-ground and above-ground nutrients in pepper plants. Understanding the intricate relationships between microbial communities and nutrient cycling in the rhizosphere is crucial for optimizing agricultural practices and enhancing crop productivity while reducing reliance on chemical fertilizers and pesticides.

In certain tissues of pepper plants, the levels of TN and TP nutrients were consistently lower in LN and HR varieties compared to the other two peppers (Fig. 1). This can be attributed to the presence of common associated microbes and their functional genes, particularly those involved in N/P cycling processes. The analysis revealed that the abundance of N/P-associated genes was closely correlated with bacterial network clusters in module #8 and module #15, as well as the changes in TN and TP content in different parts of the pepper plants. Interestingly, certain microbial genera, such as *Jatrophihabitans*, *Conexibacter* (Wu *et al*., 2022), *Rhodanobacter* (Wang *et al*., 2022; Wu *et al*., 2022), f_*67-14*, *Acidibacter* (Song *et al*., 2022), f_*Microscillaceae*, and f_*Chitinophagaceae*, were found to be enriched in specific pepper varieties and likely play significant roles in N/P cycling. On the other hand, high-abundance genera in YZ and BE cultivars, including *Arthrobacter*, *Nitrosospira*, *Frankiales*, *Bacillus*, *Lysobacter*, *Sorangium*, *Conexibacter*, and *Leifsonia*, have been reported to be involved in nitrification or nitrogen fixation processes (Hu *et al*., 2021; Sellstedt and Richau, 2013; Wen *et al*., 2022; Yu *et al*., 2022). In contrast, LN and HR varieties showed an abundance of genera such as *Rhodanobacter*, *Pseudomonas*, *Saccharimonadales*, *Steroidobacter*, and *Rhizobiaceae*, which are associated with denitrification and ammonium oxidation processes (Hou *et al*., 2018; Wang *et al*., 2022; Zhu *et al*., 2023). These functional genes likely play a crucial role in the soil rhizosphere, directly impacting nutrient cycling and indirectly influencing the nutritional traits of the pepper plants. Interestingly, a strong positive correlation was observed between the abundance of CNPS cycling genes and the nutrient content in pepper plant parts, rather than with soil nutrient properties. This suggests the presence of functional redundancy in the soil microbiome (Xiang *et al*., 2020) or the influence of anthropogenic factors such as fertilization practices.

It was challenging to quantitatively measure the specific effects of individual microorganisms or functional genes on plant physiological processes. However, in this study under our controlled conditions, we observed a close relationship between these genes and the nutrient properties of pepper plants. The distinct taxa and their differentially expressed genes in the rhizosphere explained a significant portion of the variation in below- and aboveground nutrient levels of the pepper plant. Through structural equation model analysis, we considered factors such as microbial modules, functional genes, soil physicochemical properties, and below/aboveground nutrient properties of peppers. The analysis revealed that the total effects on aboveground nutrients were influenced in the order of root > microbes > microbial genes > soil properties, explaining a high variance (R^2^=92%) in the aboveground nutrient features of pepper plants. In conclusion, our research, based on statistical correlation analysis, provides valuable insights into understanding the effects of rhizospheric microbes and functional genes on pepper nutrient properties (both below- and aboveground) under controlled experimental conditions. However, it is important to note that the C/N/P/S metabolic genes detected by the q-PCR chip did not always exhibit the same changes as the nutrient variation in pepper plants. This discrepancy could be attributed to hysteresis effects occurring over different time and spatial scales. Further research should focus on studying the nutrient accumulation patterns throughout the developmental stages of peppers and the corresponding microbiota at different periods. This will help us gain a better understanding of the specific functions of microbes in the pepper rhizosphere.

## Conclusion

Our findings indicate that the soil rhizosphere of pepper cultivars possesses a diverse and significant microbial community, along with a wide range of CNPS cycling genes. These factors play a crucial role in shaping the distribution patterns of microbial communities and their associated CNPS cycling in rhizosphere soils. Through network analysis and correlation calculations, we discovered that the bacterial community exhibited more abundant associations compared to the fungal community. Furthermore, our structural equation models effectively explained the strong relationships between microbial modularity, functional genes, and below-/above-ground nutrient traits in pepper plants. However, to gain a deeper understanding of the effect patterns of microbial clusters and individual functional genes on pepper nutrient accumulation, further controlled experiments are necessary.

## AUTHOR CONTRIBUTIONS

Li X. and Tao Y.: Conceptualization. Peng SW.: Investigation, Methodology; Zhang MX.: Visualization, Writing-original Draft. Tao Y.: Review & Editing; Zou XX. and Long SP.: Funding acquisition. All authors contributed to revisions and gave final approval for publication.

## ACKNOWLEDGMENTS

This work was supported by the Science and Technology Project of Hunan Province (2021NK1040); Agricultural Science and Technology Innovation of Hunan Province (2022CX39); and Key R&D Program of Hunan Province (2021NK2006); Hunan Young Science and technology Talent Project (2022RC1047).

## DATA AVAILABILITY

The raw sequences of 16S rRNA genes and ITS regions described in this article have been submitted to the NCBI Sequence Read Archive (SRA) database (accession number PRJNA981279).

